# Histone methylation activity of KMT2D is required for proliferative control of the developing lung

**DOI:** 10.64898/2026.03.18.712755

**Authors:** Aalekh S. Mehta, Guojia Xie, Frances A. High, Patricia K. Donahoe, Samuel P. Rowbotham

## Abstract

KMT2D, the histone methyltransferase and core component of the COMPASS/MLL4 complex, has been implicated in developmental diseases such as Kabuki Syndrome, interstitial lung disease, and congenital diaphragmatic hernia, with clear links to pediatric pulmonary disorders. Despite this, the mechanism by which KMT2D governs lung development remains unclear. Knock-in mouse models rendering, KMT2D catalytically deactivated (KMT2D^KI^) and reducing H3K4 methylation, have demonstrated potential in defining KMT2D’s role in pulmonary development. Our examination of the lungs of KMT2D^KI^ mice revealed increased cellular density and impaired sacculation indicated by reduced airspace chord length, thickening of intersaccular septa, and abnormal alveolar cell differentiation. KMT2D^KI^ mice revealed narrowed Sox2+ conducting airways and epithelial differentiation defects characterized by reduced Cc10+ club cells. Accompanying the alveolar and airway hypoplasia, blood vessel luminal area was reduced. Conversely, KMT2D^KI^ lungs had a significantly higher proportion of proliferating cells accompanied by a dramatic expansion in Pdgfrα+ mesenchymal progenitor cells. Our findings therefore suggest that KMT2D-mediated H3K4 methylation is vital to normal lung development, and its impairment results in widespread pulmonary hypoplasia and potentially pulmonary hypertension.

## Introduction

The packaging of DNA into chromatin allows for the compaction of linear DNA into the nucleus, where it regulates gene expression [1]. Making up the structure of chromatin is the histone octamer, the eight-protein complex constituting the nucleosome. The nucleosome contains paired copies of 4 core proteins, H2A, H2B, H3, and H4 [1].

Histones are post-translationally modified to regulate gene expression during normal development and changes in homeostasis [1]. These post-translational modifications—methylation, acetylation, ubiquitination, and phosphorylation—are guided by specialized enzymes [2]. Histone methyltransferases are the enzymes responsible for methylation of these histones, and this can entail mono-, di-, or tri-methylation of amino acid residues [3]. In the context of H3, methylations of lysine at amino acid positions 4, 9, 36, and 79 (H3K4, H3K9, H3K36, and H3K79, respectively) exhibit a critical role in embryonic development [4]. Although there is an abundance of literature characterizing the role of H3K4 methylation in cardiac diseases [5], neurodevelopmental defects [6], cancers [7], and metabolic diseases [8], much remains to be explored in lung pathology and development. Modifications of H3K4 are largely carried out by the COMPASS Complex (COMplex of Proteins ASsociated with Set1), a family of methyltransferases and other subunit modifiers that work to regulate gene expression via modulation of enhancers, promoters, and gene bodies [9]. The KMT2D/MLL4 COMPASS-driven modifications of H3K4me1 and H3K27ac together are key signatures of active enhancers and ultimately facilitate development through regulation of gene expression [10]. Pathogenic variants in *KMT2D* (Lysine-specific methyltransferase 2D), the gene encoding the histone methyltransferase primarily responsible for H3K4 mono-methylation (H3K4me1) and di-methylation (H3K4me2) at enhancers, cause Kabuki Syndrome, a disease largely characterized by craniofacial abnormalities and developmental delays [11, 12]. Additionally, emerging case reports have identified a subset of patients with pathogenic *KMT2D* variants exhibiting pediatric pulmonary hypertension, pulmonary hypoplasia, and interstitial lung disease [13, 14, 15, 16].

Similarly, we have shown previously in genomic analyses of large patient cohorts that pathogenic variants in *KMT2D* play a role in the development of human Congenital Diaphragmatic Hernia (CDH), a birth defect in which patients exhibit severe lung hypoplasia characterized by neonatal respiratory distress and a high mortality rate [17, 18]. Additionally, in lung cancers *KMT2D* typically acts as a tumor suppressor [19], and a reduction in its expression facilitates the progression of lung squamous cell carcinoma [20, 21, 22]. Despite this pivotal clinical significance in pulmonology, little is known about *KMT2D*’s function in normal embryonic lung development.

As KMT2D is the key scaffold protein that forms the core of COMPASS Complex, it is required to maintain the function of the complex subunits [23]. Without KMT2D, COMPASS subunits such as KDM6A (Lysine-specific demethylase 6A), the H3K27me demethylase, show reductions in protein levels [24, 25], and other members such as PTIP (PAX transcription activation domain interacting protein) and PA1 (PAX-interacting protein1-associated glutamate rich protein 1), both with key roles in transcriptional regulation and development [26, 27], lose biochemical association with COMPASS components [28]. Therefore, functional analysis of KMT2D using a complete knockout model is difficult as it would influence other COMPASS members. Recent investigation into KMT2D’s role in development using this complete knockout mouse model resulted in embryonic death at E9.5 through prevention of gastrulation, indicating vital function in early embryonic development but precluding study of later influences on the lung [11].

However, a novel model selectively inhibiting KMT2D’s enzymatic activity of H3K4me1/2 was recently developed [29]. In this model, a point mutation was applied to KMT2D’s SET domain, causing catalytic deactivation. Consequently, the effects on other members of the COMPASS complex were minimized, and the model led to later perinatal lethality with lung hypoplasia, which allowed for a specific functional analysis of KMT2D in lung development. Given not only that mutations in *KMT2D* are potential key contributors to the pulmonary pathologies found in Kabuki Syndrome [13, 14, 15, 16], CDH [17, 18, 30], and various lung cancers [19, 20, 21, 22], but also that this catalytic deactivation model’s lung phenotype remains largely uncharacterized, we used this construct to investigate the role of KMT2D in embryonic lung development. Mice with catalytically deactivated KMT2D (KMT2D^KI^) at E18.5 exhibited an increased cellular density that correlated with the severe hypoplasia in the lung, impaired sacculation, defects in airway differentiation and remodeling, and a hyperproliferation of the mesenchyme.

## Materials and Methods

### Mice

KMT2D Y5477A knock-in mouse lines were generated using CRISPR/Cas9 as described previously [29]. Mice were housed in microisolator cages with controlled temperature of 22 °C and humidity of 45–65%. Mice were on a 12-h light and 12-h dark cycle, with water and food ad libitum. All mouse experiments were performed in accordance with ARRIVE and other relevant guidelines and regulations, and according to protocols approved by the Animal Care and Use Committee of NIDDK, NIH (protocol K165-LERB-20).

### Histology

E18.5 embryos were delivered by cesarean section, genotyped, and fixed in 4% paraformaldehyde overnight at 4°C before being dehydrated and embedded in paraffin.

Blocks were sectioned at 5 µm and the tissue was dried and adhered to microscope slides. Slides were left to dry overnight at room temperature before staining.

Hematoxylin and Eosin (H&E) staining was carried out using reagents from the Vector Laboratories H&E Staining Kit (Vector Laboratories Cat#H-3502) and their protocol. Microscope slides were deparaffinized with 3 washes of xylene before progressive rehydration with ethanol dilutions of 100% (3 minutes), 90% (1 minute), 70% (1 minute), and 30% ethanol (1 minute), before washing with distilled water (2 minutes). Tissue sections were completely covered in hematoxylin and incubated for 5 minutes before rinsing. Sections were then completely covered in Bluing Reagent and incubated for 10-15 seconds. Slides were dipped in 100% ethanol before application of Eosin Y Solution, followed by a 3 minute incubation. Slides were rinsed, fully dehydrated, cleared of excess ethanol, and cover-slipped using mounting medium.

### Immunohistochemistry

Immuno-staining was performed as previously described [31]. Sections were deparaffinized using dewaxing and rehydration steps. Antigen retrieval was performed using citrate-based unmasking buffer with a pH of 6 (Vector Laboratories Cat# H-3300-250). The slides were then incubated for 1 hr in 10% Normal Goat Serum (ThermoFisher Cat# 50062Z) in a humid chamber. Antibodies were applied to slides at established concentrations (Cell Signaling Anti-H3K4me1, 1:150, Cat# 5326; Abcam Anti-Podoplanin, 1:200, Cat# ab11936; Abcam Anti-Prosurfactant Protein C (proSP-C), 1:400, Cat# ab211326; Santa Cruz Anti-Hop E1 (Hopx), 1:125 Cat# sc-398703; Invitrogen Anti-Sox2 1:100 Cat# 14-9811-82; Santa Cruz Anti-CC10, 1:100, Cat# sc-390313; Sigma-Aldrich Anti-Acetylated Tubulin, 1:1000, Cat# T6793-100UL; Invitrogen Anti-Alpha-Smooth Muscle (α-SMA), 1:200, Cat# 14-9760-82; Santa Cruz Anti-Pdgfrα C-20, 1:215, Cat# sc-338; Invitrogen Anti-KI67, 1:100, Cat# 14-5698-82; Agilent Anti-Von Willebrand Factor (VWF), 1:100, Cat# A008229-5) and incubated overnight at 4C.

Slides were then washed and stained with DAPI (1:200, ThermoFisher Cat# 62248) and specific secondary antibodies (Invitrogen Goat anti-Mouse 488, 1:200, Cat# A11001; Abcam Goat anti-Mouse 647, 1:200, Cat# ab150115; Invitrogen Goat anti-Rabbit 488, 1:200, Cat# A11008; Invitrogen Goat anti-Rabbit 647, 1:200, Cat# A21245; Invitrogen Goat anti-Rat 488, 1:200, Cat# A11006; Invitrogen Goat anti-Hamster 647, 1:200, Cat# A21451) for an hour at room temperature. The tissue was washed and received a drop of mounting medium before coverslipping. Images were acquired via a Nikon Eclipse fluorescence microscope with a 4x, 10x, 20x, and 40x objectives, using NIS-Elements software (NIS-Elements BR 5.02.01). Images were processed uniformly using NIS-Elements with identical exposure across experimental groups.

### Morphometric Analysis

Sections underwent morphometric analysis via microscopy tools that measure distance relative to pixel size. Built-in NIS Elements software generated values measuring airspace chord length and septal wall thickness. Luminal area was measured using the trace measurement function. Blood vessels were identified by Von Willebrand Factor (VWF+) and conducting airways by Sox2+ staining. Vascular wall thickness and airway smooth muscle thickness was quantified by measuring the thickness of the α-SMA staining in VWF+ blood vessels and Sox2+ conducting airways, respectively. The Mean Linear Intercept (MLI) of lungs was calculated using a 12×8, 50-μm grid in accordance with standard protocols [32].

### Statistics and Reproducibility

Data expressed in figures are mean ± SD. Both experimental groups contained three biological replicates (n=3 mice per genotype). To reduce sources of variation such as staining batch, developmental timing, and maternal environment, each wild-type mouse had a corresponding litter-matched mutant, and staining for each marker was performed on all 6 mice in simultaneous batches. Thus, all statistical tests carried out were two-tailed unpaired t-tests with Welch’s correction, which account for any potential inherent differences in the standard deviations between the two populations, and further reduces Type 1 error rate. Positive cells were quantified as percentage of DAPI+ nuclei per field, or as percentage of DAPI+ nuclei per airway, depending on marker. GraphPad Prism 10 was used to conduct statistical tests, log data, and generate graphs. Quantification was performed using predefined criteria; for each mouse, nonadjacent sections were analyzed, and 5 random fields per section were quantified. The significance level was set to α = 0.05.

## Results

### KMT2D^KI^ Mice Exhibit Dense and Hypoplastic Lungs

We first confirmed the lack of enzymatic function in the KMT2D^KI^ lungs by quantifying the staining intensity of H3K4me1, one of the primary histone modifications established by KMT2D, in E18.5 wild type and KMT2D^KI^ embryos. The lungs were found to have significantly reduced H3K4me1, demonstrating a major loss of KMT2D methylation activity (Figure 1A). Staining of other organs revealed similar reductions in H3K4me1 (Figure 1B). We then stained embryo sections to assess general lung and thoracic cavity morphology and confirmed the earlier examination that KMT2D^KI^ embryos exhibited hypoplastic lungs^31^, though diaphragms remained intact without any apparent herniation (Figure 1C). The average area of the lungs was not significantly different between KMT2D^KI^ and wild type embryos; however, the thoracic cavity area and inter-thoracic space trended towards smaller areas in KMT2D^KI^ embryos (Figure 1D). Both DAPI+ and Hematoxylin-dyed nuclei were counted and shown to be increased in KMT2D^KI^ mice compared to those of the wild type, indicating excessive cell density (Figure 1E). Taken together, the knock-in deactivation of KMT2D demonstrated dense and morphologically abnormal lungs at E18.5 in KMT2D^KI^ compared to wild type mice.

**Figure 1.**
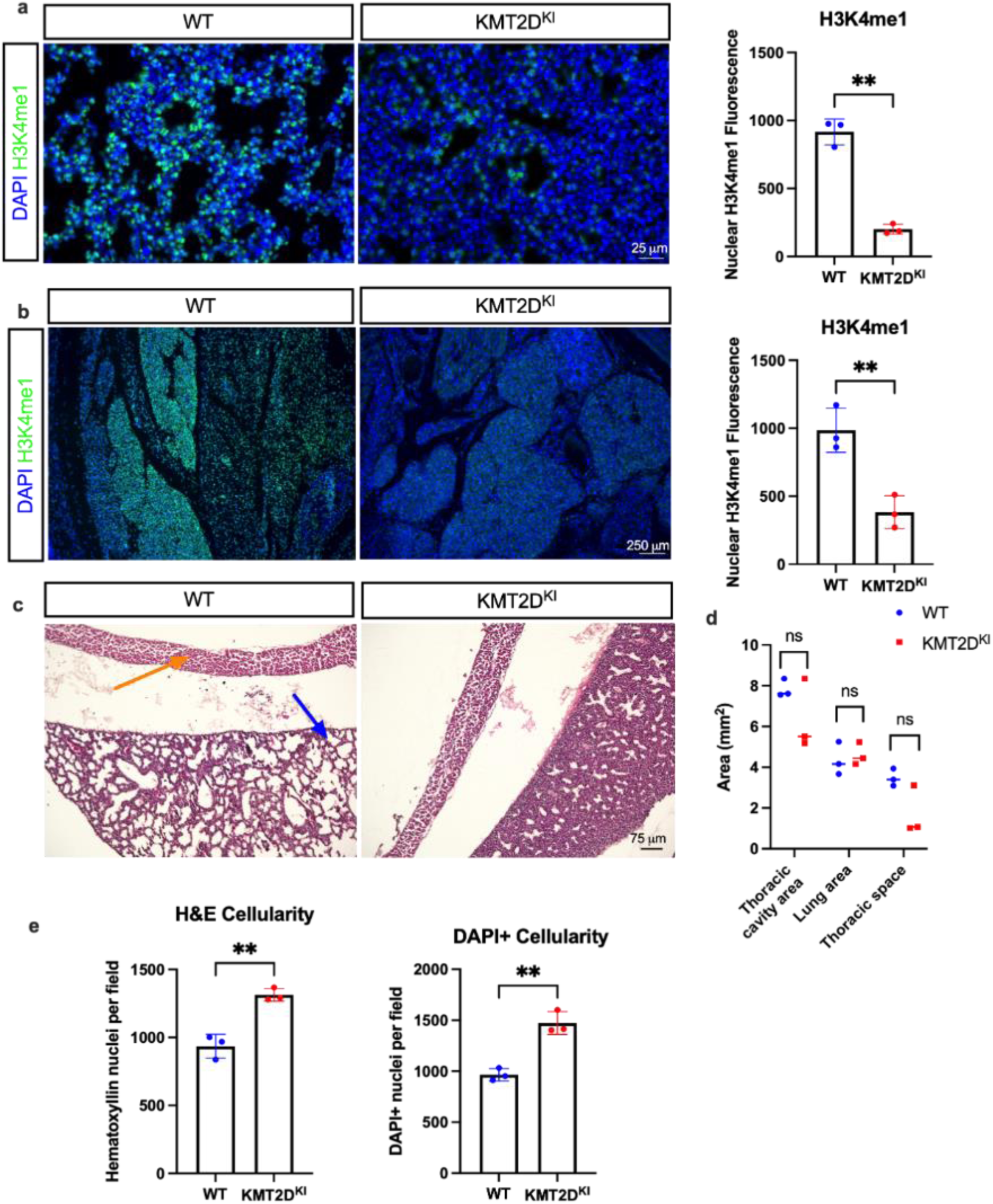
KMT2D^KI^ mouse lung density is increased alongside reduction of H3K4me1. A) Immunostainings of H3K4me1 in E18.5 WT and KMT2D^KI^ lungs with quantification (p=0.0025). Scale bar = 25µm. B) Immunostaining of H3K4me1 in gastrointestinal organs of E18.5 WT and KMT2D^KI^ mice (p=0.0084). Scale bar = 250µm. C) H&E staining of lungs and diaphragms of E18.5 WT and KMT2D^KI^ mice. Orange arrow indicates diaphragm. Blue arrow indicates lung. Scale bar = 75µm. D) Quantification of thoracic cavity area (p=0.2732), lung area (p=0.6809), and thoracic space (p=0.1144) in WT and KMT2D^KI^ mice. E) Quantification of E18.5 DAPI+ (p=0.0057) and hematoxylin+ nuclei (p=0.0064) in lungs of WT and KMT2D^KI^ mice. *=p<0.05, **=p<0.01 by Welch’s two-tailed unpaired t-test. Graphs depict means ± SD.

### Impaired distal lung sacculation

H&E staining of wild type and KMT2D^KI^ mice was further analyzed to assess lung morphometrics. Saccular chord length was significantly reduced in mutant mice compared to controls, implying underdevelopment of the future alveoli (Figure 2A,B). Average saccule septal wall thickness was measured and found to be thicker in mutants, suggesting diminished effective gas exchange (Figure 2A,C). Mean linear intercept (MLI) measurements demonstrated miniaturized saccules in mutant mice, confirming these findings (Figure 2A,D).

**Figure 2.**
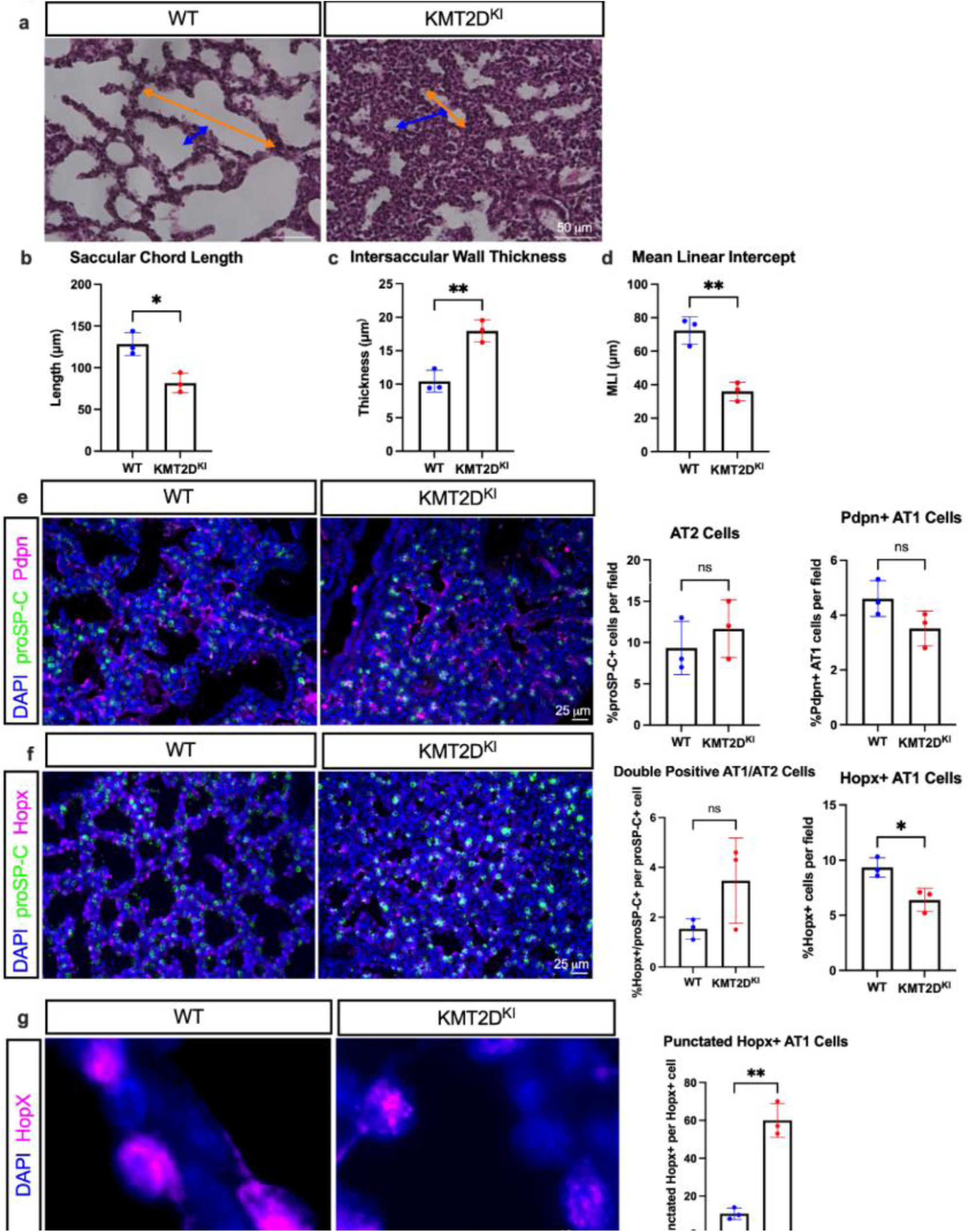
KMT2D^KI^ mice exhibit underdeveloped distal lungs and a Hopx+ AT1 cell deficit. A) H&E staining of WT and KMT2D^KI^ mouse lung. Orange arrow indicates measurement of pre-alveolar saccular chord length. Blue arrow indicates measurement of intersaccular septal wall thickness. Scale bar = 50µm. B) Quantification of saccular chord length (p=0.0119). C) Quantification of intersaccular septal wall thickness (p=0.0050). D) Quantification of mean linear intercept (p=0.0047). E) Immunostaining and quantification of proSP-C+ AT2 (p=0.4441) and Pdpn+ AT1 (p=0.1074) alveolar cells in WT and KMT2D^KI^ lungs. Scale bar = 25µm. F) Immunostaining and quantification of Hopx+ AT1 population (p=0.0214) and proSP-C+/Hopx+ transitional AT1/AT2 cells (p=0.1840) in WT and KMT2D^KI^ lungs. Scale bar = 25µm. G) Comparative immunostaining and quantification of diffuse and punctated Hopx+ AT1 cells in WT and KMT2D^KI^ lungs (p=0.0059). Scale bar = 10 µm. *=p<0.05, **=p<0.01 by Welch’s two-tailed unpaired t-test. Graphs depict means ± SD.

To dissect the abnormal saccular structure further, we stained for markers of alveolar type I (AT1) and II (AT2) cells, responsible for carrying out gas exchange and surfactant production. Immunostaining of AT2 marker Pro-Surfactant Protein C (proSP-C) found that the AT2 cell population was not significantly altered, despite significant differences in saccule architecture (Figure 2E). Staining of AT1 markers Podoplanin (Pdpn) and Hopx indicated a slight reduction in AT1 cell abundance, reaching statistical significance for Hopx but not Pdpn (Figure 2E,F). Dual staining of proSP-C and Hopx suggested a possible increase in transitional ATI/ATII population in KMT2D^KI^ mice but also did not reach statistical significance (Figure 2F). Coinciding with the difference observed in the number of Hopx+ cells, KMT2D^KI^ mutants exhibited a distinct pattern of punctate staining within the nuclei of AT1 cells, as opposed to the diffuse nuclear staining found in the wild type mice (Figure 2G). This unusual distribution of HopX may be indicative of further abnormal maturation of AT1 cells.

### Airway Remodeling and Differentiation Defect

To assess the function of KMT2D in the airway epithelium, Sox2 staining was carried out to assess conducting airway morphology in WT and KMT2D^KI^ mice (Figure 3A). Morphometric analysis of the Sox2+ stained airways demonstrated a significantly reduced cross-sectional luminal surface area in KMT2D^KI^ mice, consistent with airway narrowing. To investigate these perturbed airways further, the differentiation status of the airway epithelium was probed. Immunostaining of Cc10, marking the club cell population, indicated a significant reduction in Cc10+ club cells in KMT2D^KI^ lungs compared to controls (Figure 3B). The other predominant epithelial cell type of the distal airways, ciliated cells, exhibited no difference in their population when immunostained for acetylated α-tubulin (Figure 3C). Furthermore, staining of alpha smooth muscle actin (α-SMA) revealed no differences in peribronchiolar wall thickness between mutant and control mice (Figure 3D).

**Figure 3.**
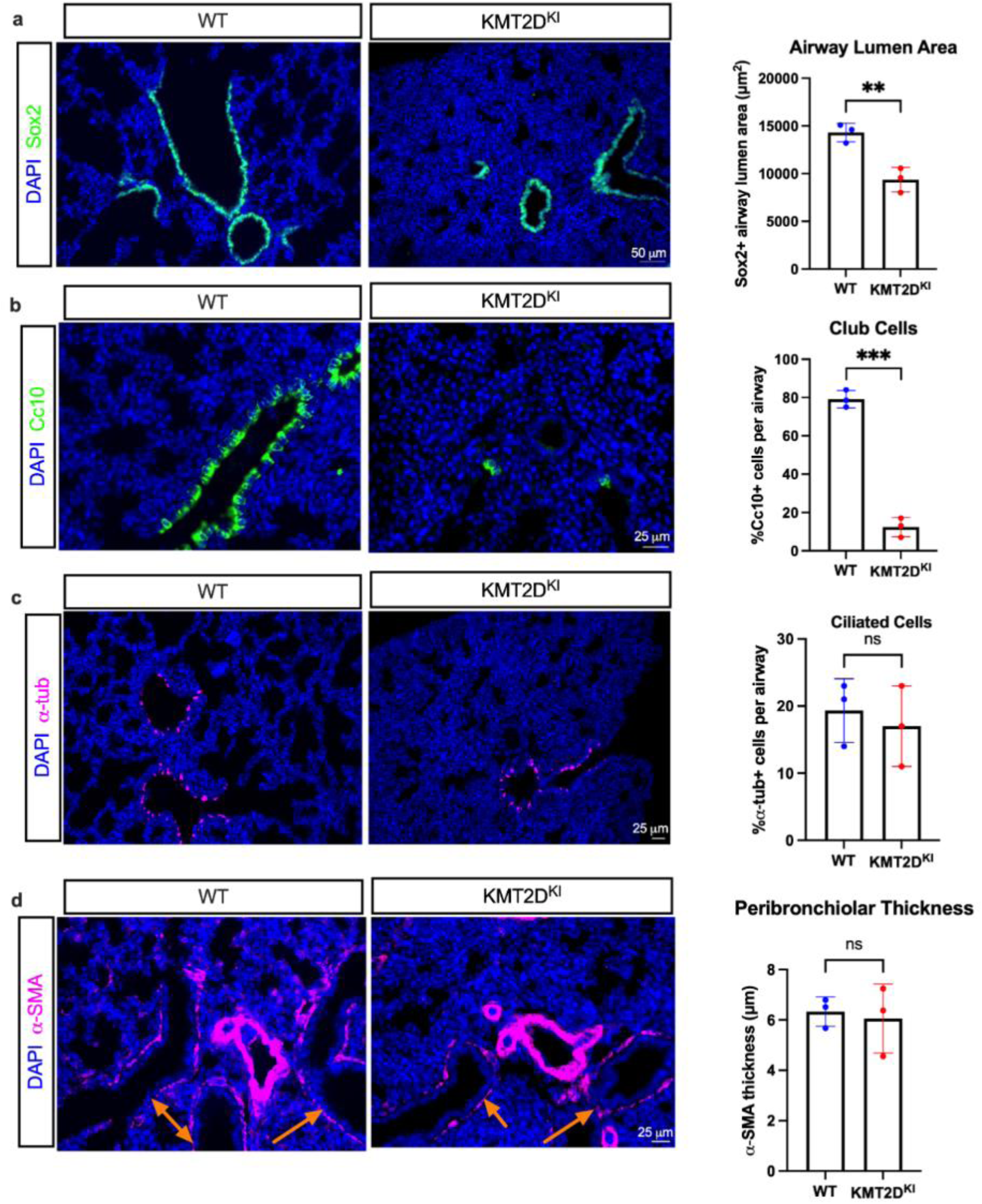
KMT2D^KI^ mice exhibit airway remodeling and differentiation defects. A) Immunostaining of Sox2+ conducting airway epithelia with quantification of enveloped airway luminal area between KMT2D^KI^ and WT mice (p=0.0074). Scale bar = 50µm. B) Immunostaining of Cc10+ club cells with quantification between KMT2D^KI^ and WT mice (p<0.0001). Scale bar = 25µm. C) Immunostaining of α-tubulin+ ciliated cells with quantification between KMT2D^KI^ and WT mice (p=0.6262). Scale bar = 25µm. D) Immunostaining of α-SMA+ smooth muscle cells with quantification of airway smooth muscle thickness between KMT2D^KI^ and WT mice (p=0.7732). Orange arrows indicate airway smooth muscle. Scale bar = 25µm. *=p<0.05, **=p<0.01, ***=p<0.001 by two-tailed unpaired t-test. Graphs depict means ± SD.

### Mesenchymal Hyperproliferation and Vascular Remodeling

To investigate differences in other cell lineages, further immunostaining of proliferative and non-epithelial markers was carried out. Immunostaining of the mesenchymal marker Pdgfrα showed a large increase in the Pdgfrα+ population in KMT2D^KI^ mutant lungs (Figure 4A). As we had already noted an increased cell density in the KMT2D^KI^ lungs, we sought to test if a change in the proliferative state of the mesenchymal compartment could account for this. Immunostaining of the proliferation marker Ki67 revealed significantly increased Ki67+ cells throughout the KMT2D^KI^ lungs compared to the controls, coinciding with the expanded Pdgfrα+ compartments (Figure 4B). We did not observe a difference in α-SMA+ cells in the distal lung, suggesting there was no change in differentiation towards the secondary crest myofibroblast fate. Assessment of vascular morphology via alpha-smooth muscle actin (α-SMA) and the endothelial marker Von Willebrand Factor (VWF) showed no difference in vascular wall thickness between controls and KMT2D^KI^ mice (Figure 4C,D). However, cross-sectional vessel lumen surface area was reduced in mutant lungs (Figure 4C,E). Such a reduction in luminal area is potentially indicative of pulmonary hypertension and an additional indicator of morphological remodeling in mutant lungs.

**Figure 4.**
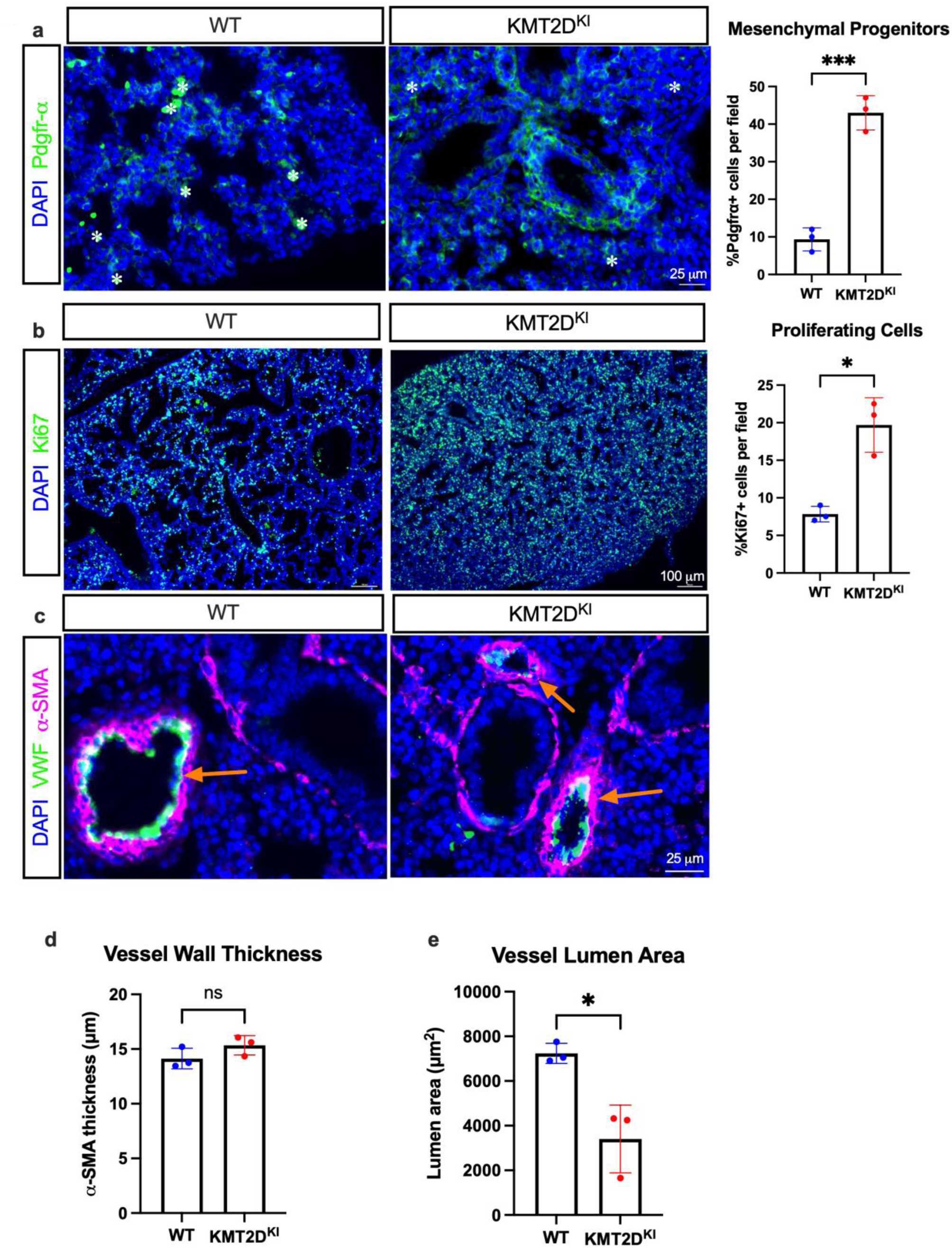
KMT2D^KI^ mice exhibit hyperproliferation alongside mesenchymal overgrowth and vascular remodeling. A) Immunostaining of Pdgfrα+ fibroblasts with quantification between KMT2D^KI^ and WT mice (p=0.0009). White asterisk = auto-fluorescent staining. Scale bar = 25µm. B) Immunostaining of Ki67+ proliferating cells with quantification between KMT2D^KI^ and WT mice (p=0.0228). Scale bar = 100µm. C) Immunostaining of α-SMA+ smooth muscle cells and VWF+ endothelial cells in WT and KMT2D^KI^ mice. Scale bar = 25µm. D) Quantification of vascular wall thickness between WT and KMT2D^KI^ mice (p=0.1808). E) Quantification of vascular luminal area between WT and KMT2D^KI^ mice (p=0.0393). Orange arrows indicate blood vessels. *=p<0.05, **=p<0.01, ***=p<0.001 by two-tailed unpaired t-test. Graphs depict means ± SD.

## Discussion

While previous literature demonstrates loss of KMT2D correlates with the progression of lung cancers [19, 20, 21, 22], can be causative for CDH [17, 18, 30], and has a demonstrated role in pediatric pulmonary disorders in Kabuki Syndrome [13,14,15,16], its role in lung development requires further investigation. The data reported here support catalytic deactivation of KMT2D causing pulmonary hypoplasia in mice. By modeling a deactivation of KMT2D’s enzymatic function while keeping the protein intact, the effects on the stability of the COMPASS Complex were minimized, allowing KMT2D-mediated H3K4me1/me2 catalyzation to be studied. As KMT2D is integral to the function of other COMPASS Complex members [24, 25, 26, 27, 28], such precision is critical to investigating its specific function in the lung.

Our data first shows that the loss of H3K4me1 in the lung causes severe pulmonary abnormalities associated with elevated cellular density, despite no obvious defects in other tissues. Although thoracic cavity area and thoracic space only trend towards significance, it is plausible that an increased sample size would yield a statistically significant reduction in both categories. Additionally, the reduction in saccular size, measured via airspace chord length, has previously been associated with impaired gas exchange [33, 34]. These findings are further corroborated by the reduced mean linear intercept (MLI) in KMT2D^KI^ mouse lungs; although an increase in MLI is the typical hallmark of several pulmonary diseases [35], MLI is generally inversely related to cellular density [36], thus our reduced MLI validates our findings of hypercellularity. The intersaccular septal thickening increases the diffusion distance between the future alveoli and reduces the Diffusing Capacity of the Lungs for Carbon Monoxide (DLCO), a common marker of pulmonary emphysema, dyspnea, fibrosis, and reduced gas exchange [37]. Notably, airspace wall thickening has been reported in patients with Kabuki Syndrome [14, 38]. This thickening, coupled with the increased cellular density, suggests that KMT2D’s function in cellular proliferation, possibly related to its tumor suppressor function, which may contribute to the eventual structural abnormalities exhibited in KMT2D^KI^ lungs, as this increased density may be compressing the alveoli. Further, lungs in patients with Kabuki Syndrome have exhibited diffuse ground-glass opacities [39], which can be indicative of increased interstitial density and fibrosis [40]. Patients with Kabuki Syndrome with pulmonary disorders have demonstrated diffuse alveolar hemorrhage [41], and these alveolar defects in the mutant mice may explain its functional severity. Alongside this clear impairment in sacculation, the population of AT1 cells was also reduced. Additionally, staining of the key AT1 transcriptional factor Hopx revealed a difference in nuclear localization, further raising the possibility of a defect in AT1 cell differentiation. Reduced Hopx expression in the lung has been implicated as a key signature of declined lung function and idiopathic pulmonary fibrosis [42]. Relatedly, this reduction in Hopx expression might reflect a specific interaction between H3K4me1, KMT2D-COMPASS, and Hopx. Mechanistically, Hopx promoters have been shown in bone marrow stromal cell differentiation to interact with the H3K27 methyltransferase EZH2 (Enhancer of Zeste Homolog 2), an enzyme that opposes the function of demethylase KDM6A, a COMPASS subunit that associates with KMT2D [43]. Taken together, the impaired sacculation and abnormal Hopx expression may be indicative of disrupted epigenetic regulation as measured by the reduction of H3K4me1.

Importantly, a recent update on KMT2C [44], a paralog of KMT2D, in lung development showed that a conventional knockout of KMT2C generated lungs developmentally similar to those of our KMT2D model. Lungs appeared similarly dense, with a near total reduction of Cc10+ club cells. The predominant finding in this KMT2C model was a widespread AT1 defect, in which staining of AT1 marker Aqp5 demonstrated KMT2C mutant lungs as deficient in AT1 cells. Although KMT2C’s role in mesenchymal proliferation was not surveyed in this study, total proliferating cells were increased; importantly, this increase in proliferation in KMT2C mutant lungs was accompanied by an increased deposition of fibronectin and laminin, ECM proteins that are largely produced by fibroblasts and myofibroblasts [45]. Together, this is suggestive of redundancies between KMT2C and KMT2D. A notable difference and potential lack of redundancy between the two paralogs, however, is the AT1 defect: while our model showed a specific Hopx defect in the AT1 population, the KMT2C mutant AT1 defect findings were further confirmed by qRT-PCR and mRNA expression profiling of other AT1 markers including Hopx, Pdpn, and Clic5. As this is indicative of a general AT1 defect instead of a Hopx defect, this potentially implicates KMT2D and KMT2C both as key regulators of AT1 development, but with KMT2D’s role being mechanistically tied more to interactions with Hopx. This data suggests that future investigations of these paralogs’ redundancies should address AT1 cell development.

Coinciding with evident alveolar abnormalities, airways exhibited a profound club cell differentiation defect. Club cell reduction is associated with several lung diseases such as chronic obstructive pulmonary disorder (COPD) [46] and asthma [47], and is a hallmark signature of the pulmonary hypoplasia found in human CDH patient tracheal aspirates in vitro [48]. As club cells are key immunomodulators of the lung [46], such a stark reduction upon KMT2D’s deactivation may explain the increased susceptibility to respiratory infections, a noted pathology of Kabuki Syndrome [49]. Airway luminal area reduction can lead to a decreased air influx and may lead to air-trapping [50, 51], which is a documented pathology in Kabuki Syndrome patients [39]. Because airway stenosis is a noted finding in Kabuki Syndrome patients [52], further investigation into the physiological consequences of this airway remodeling effect is warranted.

Immunostaining has revealed that this increased cellular density coincides with an increased population of Ki67+ proliferating cells; however, the precise identity of these proliferating cells remains unresolved. As the Pdgfrα+ mesenchymal population was the only lineage that exhibited a significant increase, this suggests that the proliferation and increased cell density was likely mesenchyme-based, though lineage tracing would be needed to confirm this. Notably, patient-derived Kabuki Syndrome iPSCs have exhibited upregulation of mesenchymal development genes such as TBX2, TWIST1, GBX2, MEF2C, supporting the claim of mesenchymal overgrowth contributing to this lung hypoplasia in KMT2D^KI^ mice [53]. Existing research implicates KMT2D’s function in epithelial-to-mesenchymal transition and has indicated that loss of KMT2D-COMPASS pushes cells into a fully mesenchyme state, but little is known about its function in the lung mesenchyme [54]. Regardless, the effect of this hyperproliferation extended beyond epithelial and mesenchymal cells to the structure of the endothelium, as additional morphometrics of the vasculature revealed a reduction in blood vessel luminal area. As the vessel luminal area is reduced, equivalent blood flow through a smaller space can lead to pulmonary hypertension. This relationship between pulmonary vascular stenosis and hypertension has been documented in CDH patients [55]. Taken together, future investigation of conditional knockouts of KMT2D in the lung mesenchyme may be warranted to examine whether the widespread lung hypoplasia is a result of the mesenchymal overgrowth.

As KMT2D-COMPASS-driven histone modifications of H3K4me1 and H3K27ac are responsible for enhancer activity [10], modulating these functions may be fruitful in rescuing the KMT2D^KI^ lung developmental defects and eventually treating Kabuki Syndrome and CDH patients with this mutation. Recently, dexamethasone, a corticosteroid that has been shown to modulate H3K27ac and H3K27me3 [56, 57], was found to successfully both rescue the lung hypoplasia of the rat nitrofen CDH model [58] and rescue club cell differentiation defects in human CDH patient tracheal aspirate cultures [48]. Additionally, EZH2, the gene responsible for downregulating COMPASS-driven enhancer activity through H3K27me3, and its complex (PRC2) was identified as a gene with enriched targets of epigenetic regulation by bulk RNA sequencing between human CDH and control patient tracheal aspirates [48]. Because modulation of EZH2 has been shown to act as a molecular switch that determines differentiation from basal cells to secretory cells [59], and because treatment with dexamethasone indicates its role in regulating differentiation of epithelial progenitors [60], there is merit to treating KMT2D^KI^ lung explants with a drug targeting this COMPASS-opposing pathway in pursuit of a developmental rescue.

In summary, we revealed that reduction of H3K4me1 via catalytic inactivation of KMT2D has a critical effect on lung development, regulating morphology, cell proliferation, and differentiation in mouse lungs. KMT2D inactivity is further associated with reduction in luminal areas of all major parts of the lung, so further investigation into the relationship between hyperproliferation of the mesenchyme and the subsequent hypoplasia and hypertension is merited. Future studies will be needed to test whether a mesenchymal signal, either absent or overexpressed, results in hypoplasia of the epithelial, endothelial, and alveolar compartments which this model presents.

## Acknowledgments

We thank current and past members of the Pediatric Surgical Research Laboratory at Massachusetts General Hospital. We thank Kai Ge at the National Institute of Diabetes and Digestive Kidney Diseases for the generation of the KMT2D^KI^ mutant mouse model. This work was supported by the Intramural Research Program of NIDDK, (G.X.), National Institutes of Health (R01HD115718) to F.A.H., and the National Institutes of Health (1R03HD112717-01), American Lung Association Catalyst Award CA-1052191, and Charles Hood Foundation Child Health Research Award to S.P.R. The content is solely the responsibility of the authors and does not necessarily represent the official views of the National Institutes of Health.

## Author Contributions

ASM Investigation, Formal Analysis, Methodology, Conceptualization, Writing – Original Draft GX Resources, Methodology, Conceptualization, Writing – Review FAH Conceptualization, Writing – Review, Funding acquisition PKD Conceptualization, Writing – Review, Funding acquisition SPR Methodology, Conceptualization, Writing – Original Draft, Supervision, Funding acquisition

All authors approved the submitted version.

## Data availability

The data supporting the findings of this study are available from the corresponding author upon reasonable request.

## Additional Information

The authors declare no competing interests.

## References

1. Sokolova, V., Sarkar, S. & Tan, D. Histone variants and chromatin structure, update of advances. Comput. Struct. Biotechnol. J. 21, 299–311 (2022).

2. Liu, R. et al. Post-translational modifications of histones: mechanisms, biological functions, and therapeutic targets. MedComm 4, e292 (2023).

3. Husmann, D. & Gozani, O. Histone lysine methyltransferases in biology and disease. Nat. Struct. Mol. Biol. 26, 880–889 (2019).

4. Bozdemir, N. et al. A comprehensive review of histone modifications during mammalian oogenesis and early embryo development. Histochem. Cell Biol. 163, 70 (2025).

5. Zhang, Q. J. & Liu, Z. P. Histone methylations in heart development, congenital and adult heart diseases. Epigenomics 7, 321–330 (2015).

6. Vallianatos, C. N. & Iwase, S. Disrupted intricacy of histone H3K4 methylation in neurodevelopmental disorders. Epigenomics 7, 503–519 (2015).

7. Wang, H. & Helin, K. Roles of H3K4 methylation in biology and disease. Trends Cell Biol. 35, 115–128 (2025).

8. Hsu, C. L. et al. H3K4 methylation in aging and metabolism. Epigenomes 5, 14 (2021).

9. Cenik, B. K. & Shilatifard, A. COMPASS and SWI/SNF complexes in development and disease. Nat. Rev. Genet. 22, 38–58 (2021).

10. Barral, A. & Déjardin, J. The chromatin signatures of enhancers and their dynamic regulation. Nucleus 14, 2160551 (2023).

11. Lee, J. E. et al. H3K4 mono- and di-methyltransferase MLL4 is required for enhancer activation during cell differentiation. eLife 2, e01503 (2013).

12. Bögershausen, N. et al. Mutation update for Kabuki syndrome genes KMT2D and KDM6A and further delineation of X-linked Kabuki syndrome subtype 2. Hum. Mutat. 37, 847–864 (2016).

13. Deng, X. et al. Pulmonary hypertension—a novel phenotypic hypothesis of Kabuki syndrome: a case report and literature review. BMC Pediatr. 23, 429 (2023).

14. Baldridge, D. et al. Phenotypic expansion of KMT2D-related disorder: beyond Kabuki syndrome. Am. J. Med. Genet. A 182, 1053–1065 (2020).

15. Bang, T. J. et al. Pulmonary manifestations of common variable immunodeficiency. J. Thorac. Imaging 33, 377–383 (2018).

16. Ba, X. et al. Case report of Kabuki syndrome in a newborn caused by KMT2D gene mutation. Front. Pediatr. 12, 1455609 (2024).

17. Qiao, L. et al. Rare and de novo variants in 827 congenital diaphragmatic hernia probands implicate LONP1 as candidate risk gene. Am. J. Hum. Genet. 108, 1964–1980 (2021).

18. Qiao, L. et al. Common variants increase risk for congenital diaphragmatic hernia within the context of de novo variants. Am. J. Hum. Genet. 111, 2362–2381 (2024).

19. Alam, H. et al. KMT2D deficiency impairs super-enhancers to confer a glycolytic vulnerability in lung cancer. Cancer Cell 37, 599–617.e7 (2020).

20. Pan, Y. et al. KMT2D deficiency drives lung squamous cell carcinoma and hypersensitivity to RTK-RAS inhibition. Cancer Cell 41, 88–105.e8 (2023).

21. Fang, Z. et al. Somatic KMT2D loss-of-function mutations in lung squamous cell carcinoma: a single-center cohort study. J. Thorac. Dis. 16, 3338–3349 (2024).

22. Yang, Y. et al. MLL4 regulates the progression of non-small-cell lung cancer by regulating the PI3K/AKT/SOX2 axis. Cancer Res. Treat. 55, 778–803 (2023).

23. Lavery, W. J. et al. KMT2C/D COMPASS complex-associated diseases [KCDCOM-ADs]: an emerging class of congenital regulopathies. Clin. Epigenetics 12, 10 (2020).

24. Fasciani, A. et al. MLL4-associated condensates counterbalance Polycomb-mediated nuclear mechanical stress in Kabuki syndrome. Nat. Genet. 52, 1397–1411 (2020).

25. Rickels, R. et al. A small UTX stabilization domain of Trr is conserved within mammalian MLL3-4/COMPASS and is sufficient to rescue loss of viability in null animals. Genes Dev. 34, 1493–1502 (2020).

26. Lechner, M. S., Levitan, I. & Dressler, G. R. PTIP, a novel BRCT domain-containing protein interacts with Pax2 and is associated with active chromatin. Nucleic Acids Res. 28, 2741–2751 (2000).

27. Kumar, A. et al. Loss of function of mouse Pax-interacting protein 1-associated glutamate rich protein 1a (Pagr1a) leads to reduced Bmp2 expression and defects in chorion and amnion development. Dev. Dyn. 243, 937–947 (2014).

28. Shpargel, K. B. & Quickstad, G. SETting up the genome: KMT2D and KDM6A genomic function in the Kabuki syndrome craniofacial developmental disorder. Birth Defects Res. 115, 1885–1898 (2023).

29. Xie, G. et al. MLL3/MLL4 methyltransferase activities control early embryonic development and embryonic stem cell differentiation in a lineage-selective manner. Nat. Genet. 55, 693–705 (2023).

30. Scott, T. M. et al. Clinical exome sequencing data reveal high diagnostic yields for congenital diaphragmatic hernia plus (CDH+) and new phenotypic expansions involving CDH. J. Med. Genet. 59, 270–278 (2022).

31. Rowbotham, S. P. et al. Age-associated H3K9me2 loss alters the regenerative equilibrium between murine lung alveolar and bronchiolar progenitors. Dev. Cell 58, 2974–2991.e6 (2023).

32. Branchfield, K. et al. Pulmonary neuroendocrine cells function as airway sensors to control lung immune response. Science 351, 707–710 (2016).

33. Coalson, J. J. et al. Neonatal chronic lung disease in extremely immature baboons. Am. J. Respir. Crit. Care Med. 160, 1333–1346 (1999).

34. Ahlfeld, S. K. et al. Relationship of structural to functional impairment during alveolar-capillary membrane development. Am. J. Pathol. 185, 913–919 (2015).

35. Kim, W. D. et al. The association between small airway obstruction and emphysema phenotypes in COPD. Chest 131, 1372–1378 (2007).

36. Alapati, D. et al. Inhibition of β-catenin signaling improves alveolarization and reduces pulmonary hypertension in experimental bronchopulmonary dysplasia. Am. J. Respir. Cell Mol. Biol. 51, 104–113 (2014).

37. Goldin, J. & Cascella, M. Diffusing capacity of the lungs for carbon monoxide. In StatPearls (StatPearls Publishing, 2025).

38. Cappuccio, G. et al. Bronchial isomerism in a Kabuki syndrome patient with a novel mutation in MLL2 gene. BMC Med. Genet. 15, 15 (2014).

39. Leonardi, L. et al. Immune dysregulation in Kabuki syndrome: a case report of Evans syndrome and hypogammaglobulinemia. Front. Pediatr. 11, 1087002 (2023).

40. Gao, J. W. et al. Pulmonary ground-glass opacity: computed tomography features, histopathology and molecular pathology. Transl. Lung Cancer Res. 6, 68–75 (2017).

41. Glickman, S. et al. Kabuki syndrome presenting with severe anemia secondary to pulmonary hemosiderosis. Ann. Allergy Asthma Immunol. 133, S167 (2024).

42. Ota, C. et al. Dynamic expression of HOPX in alveolar epithelial cells reflects injury and repair during the progression of pulmonary fibrosis. Sci. Rep. 8, 12983 (2018).

43. Hng, C. H. et al. HOPX regulates bone marrow-derived mesenchymal stromal cell fate determination via suppression of adipogenic gene pathways. Sci. Rep. 10, 11345 (2020).

44. Ashokkumar, D. et al. MLL4 is required after implantation, whereas MLL3 becomes essential during late gestation. Development 147, dev186999 (2020).

45. Zhou, Y. et al. Extracellular matrix in lung development, homeostasis and disease. Matrix Biol. 73, 77–104 (2018).

46. Blackburn, J. B. et al. An update in club cell biology and its potential relevance to chronic obstructive pulmonary disease. Am. J. Physiol. Lung Cell Mol. Physiol. 324, L652–L665 (2023).

47. Shijubo, N. et al. Clara cell protein-positive epithelial cells are reduced in small airways of asthmatics. Am. J. Respir. Crit. Care Med. 160, 930–933 (1999).

48. Wagner, R. et al. A tracheal aspirate-derived airway basal cell model reveals a proinflammatory epithelial defect in congenital diaphragmatic hernia. Am. J. Respir. Crit. Care Med. 207, 1214–1226 (2023).

49. Comel, M. et al. Abnormal immune profile in individuals with Kabuki syndrome. J. Clin. Immunol. 45, 7 (2025).

50. McNulty, W. & Usmani, O. S. Techniques of assessing small airways dysfunction. Eur. Clin. Respir. J. 1, 25898 (2014).

51. Li, T. et al. Computed tomography-identified phenotypes of small airway obstructions in chronic obstructive pulmonary disease. Chin. Med. J. 134, 2025–2036 (2021).

52. van Haelst, M. M. et al. Unexpected life-threatening complications in Kabuki syndrome. Am. J. Med. Genet. 94, 170–173 (2000).

53. Cuvertino, S. et al. Epigenome and transcriptome changes in KMT2D-related Kabuki syndrome type 1 iPSCs, neuronal progenitors and cortical neurons. PLoS Genet. 21, e1011608 (2025).

54. Zhang, Y. et al. Genome-wide CRISPR screen identifies PRC2 and KMT2D-COMPASS as regulators of distinct EMT trajectories that contribute differentially to metastasis. Nat. Cell Biol. 24, 554–564 (2022).

55. Koyuncu, M. Z. et al. Premature monozygotic twins with congenital diaphragmatic hernia: a case report. Sudan J. Paediatr. 25, 66–70 (2025).

56. Cao, J. et al. Adrenal high-expressional CYP27A1 mediates bile acid increase and functional impairment in adult male offspring by prenatal dexamethasone exposure. Adv. Sci. 12, e2413299 (2025).

57. Misale, M. S. et al. Chromatin organization as an indicator of glucocorticoid induced natural killer cell dysfunction. Brain Behav. Immun. 67, 279–289 (2018).

58. Dylong, F. et al. Overactivated epithelial NF-κB disrupts lung development in congenital diaphragmatic hernia. Am. J. Respir. Cell Mol. Biol. 69, 545–555 (2023).

59. Snitow, M. E. et al. Ezh2 represses the basal cell lineage during lung endoderm development. Development 142, 108–117 (2015).

60. Chen, H. et al. Glucocorticoid dexamethasone regulates the differentiation of mouse conducting airway epithelial progenitor cells. Steroids 80, 44–50 (2014).

